# Polymerase pausing induced by sequence-specific RNA binding protein drives heterochromatin assembly

**DOI:** 10.1101/217968

**Authors:** Jahan-Yar Parsa, Selim Boudoukha, Jordan Burke, Christina Homer, Hiten D. Madhani

**Affiliations:** Department of Biochemistry and Biophysics, University of California, San Francisco, California 94158, USA; Chan-Zuckerberg Biohub San Francisco, CA 94158

**Author notes:** equal contributions (Selim Boudoukha, Jordan Burke).

## Abstract

Packaging of pericentromeric DNA into heterochromatin is crucial for genome stability, development and health, yet its endogenous triggers remain poorly understood^1^. A defining feature of pericentromeric heterochromatin is histone H3 lysine 9 methylation (H3K9me)^2–4^. In *S. pombe*, transcripts derived from the pericentromeric *dg* and *dh* repeat during S phase^5–7^ promote heterochromatin formation through two pathways: an RNAi-dependent mechanism involving recruitment of the Clr4 H3K9 methyltransferase complex (CLR-C) via the RITS complex^8–13^, and RNAi-independent mechanism involving an RNAPII-associated RNA-binding protein Seb1, the repressor complex SHREC, and RNA processing activities^14–19^. We show here that Seb1 promotes long-lived RNAPII pausing. Pause sites associated with sequence-specific Seb1 RNA binding events are significantly enriched in pericentromeric repeat regions and their presence correlates with the heterochromatin-triggering activities of the corresponding *dg* and *dh* DNA fragments. Remarkably, globally increasing RNAPII stalling by other means induces the formation of novel large ectopic heterochromatin domains. Such ectopic heterochromatin occurs even in cells lacking functional RITS, demonstrating that RNAPII pausing can be sufficient to trigger *de novo* heterochromatin independently of RNAi. These results uncover Seb1-mediated polymerase stalling as a new signal for nucleating heterochromatin assembly in repetitive DNA.

---

To understand how Seb1 interfaces with the transcription of *dg* and *dh* repeats to promote heterochromatin, we employed a previously identified viable, heterochromatin-defective allele, *seb1-1*^15^. When combined with mutants in the RNAi machinery, *seb1-1* eliminates pericentromeric heterochromatin, while the corresponding single mutants decrease H3K9me, indicative of partially redundant pathways^15^. We examined transcription of heterochromatin at single-nucleotide resolution and tested the impact of the *seb1-1* allele using Nascent Elongating Transcript Sequencing (NET-Seq)^20^. To analyze the intrinsic transcriptional properties of heterochromatic sequences prior to the establishment of heterochromatin assembly, we used the *clr4*∆ mutant, which lacks H3K9me and displays full derepression of most silenced chromatin regions. We compared this strain to a *clr4*∆ *seb1-1* double mutant to assess the impact of *seb1-1*. We first examined the effect of *seb1-1* on transcription of non-heterochromatic regions (Figure 1). Initial inspection revealed numerous genes with a decreased peak density at 5’ regions in the double mutant with increased peak density upstream of annotated cleavage-polyadenylation sites (often called Transcription end sites or TESs) (see Figure 1a-b for examples). To analyze these trends genome-wide, travelling ratios were computed on replicate data to assess relative polymerase progression for the 500 bp segment downstream of the transcription start site (TSS; 5’ traveling ratio) and for the 500 bp segment upstream of annotated TES (3’ traveling ratio) (Figure 1c; see Methods). A lower travelling ratio in mutant vs. wild-type implies lower pausing over the region examined in the mutant and vice-versa for higher ratios. Iterative K-means clustering revealed three groups (Figure 1d) two of which, representing 77% of genes in our dataset, are significantly impacted by the *seb1-1* allele (Figure 1d Clusters I and II; Extended Data Figure 1). The *seb1-1* mutation causes a reduced median 5’ traveling ratio and an increased median 3’ traveling ratio for both clusters (Figure 1d -- clusters I and II; Figure 1e – top and middle panels) while no significant changes were observed for the third cluster (Figure 1e – Cluster III, bottom panel; see Extended Data Figure 1 for p values). These data indicate that the *seb1-1* allele leads to decreased RNAPII pausing at gene 5’ ends with an associated increased 3’ signal; the latter may be due to polymerase release from upstream pauses.

**Figure 1.**
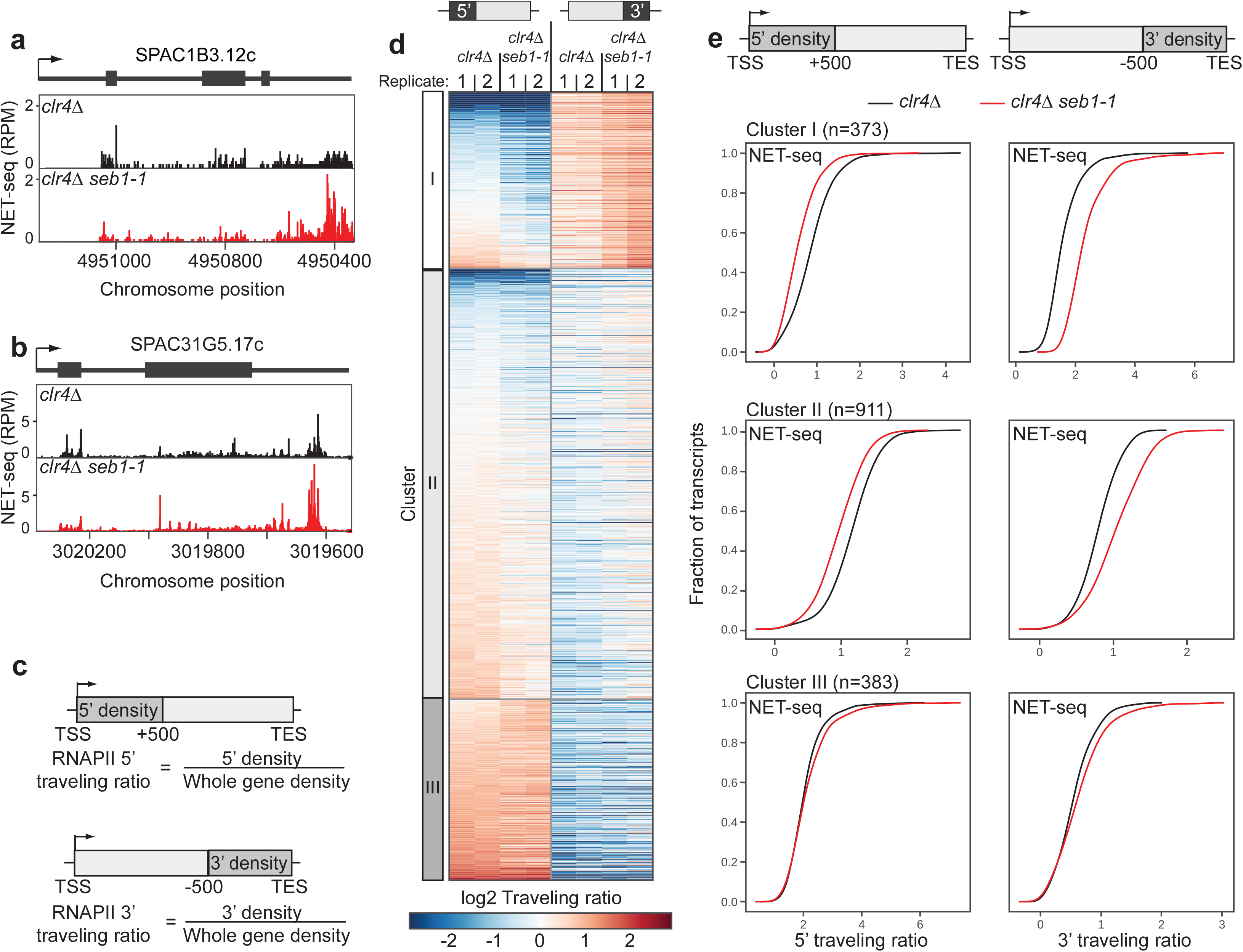
Seb1 controls polymerase progression. **a-b,** NET-seq signatures for *clr4*∆ (black) and *clr4*∆ *seb1-1* (red) strains for two genes (SPAC1B3.12c and SPAC31G5.17c). **c**, Traveling ratios at the 5’ and 3’ regions of genes. **d**, K-means clustering of NET-seq 5’ and 3’ traveling ratios (Cluster I, n=373; Cluster II, n=911; Cluster III, n=383). NET-seq replicates are represented for each *clr4*∆ and *clr4*∆ *seb1-1*. **e**, Cumulative distribution function (cdf) plots of 5’ (left column) and 3’ (right column) traveling ratios for each cluster from (**d**) comparing *clr4*∆ (black) to *clr4*∆ *seb1-1* (red) strains in each plot. KS tests were conducted for p-values (**Extended Data Figure 1**).

Our prior RIP-qPCR analysis indicates that Seb1 functions directly in heterochromatin assembly by binding pericentromeric *dg* and *dh* repeat transcripts^15^. Further, comparison of the transcriptomes of WT and *seb1-1* using RNA-seq revealed no significant changes (p<0.01 and |log_2_(fold change)| >1) in the transcript levels of known silencing factors (Extended Data Figure 2 and Extended Data Table 1). To assess direct interactions of Seb1 with pericentromeric RNA at single nucleotide resolution and across the entirety of the *dg* and *dh* regions, we conducted PAR-CLIP in replicate on a *clr4*∆ strain. Computational analysis identified statistically significant Seb1 PAR-CLIP read clusters^21^ and confirmed direct binding of Seb1 to pericentromeric transcripts (Figure 2a and b – top panels; Extended Data Figure 3a-c) via a motif described previously by others for non-heterochromatic sites bound by Seb1^14,19^ (UGUA; DREME motif analysis^22^; p=2.9e-9; E=7.4e-7). Our analysis of Seb1 PAR-CLIP read clusters for coding gene recapitulates published work is not discussed here further^14,19^.

**Figure 2.**
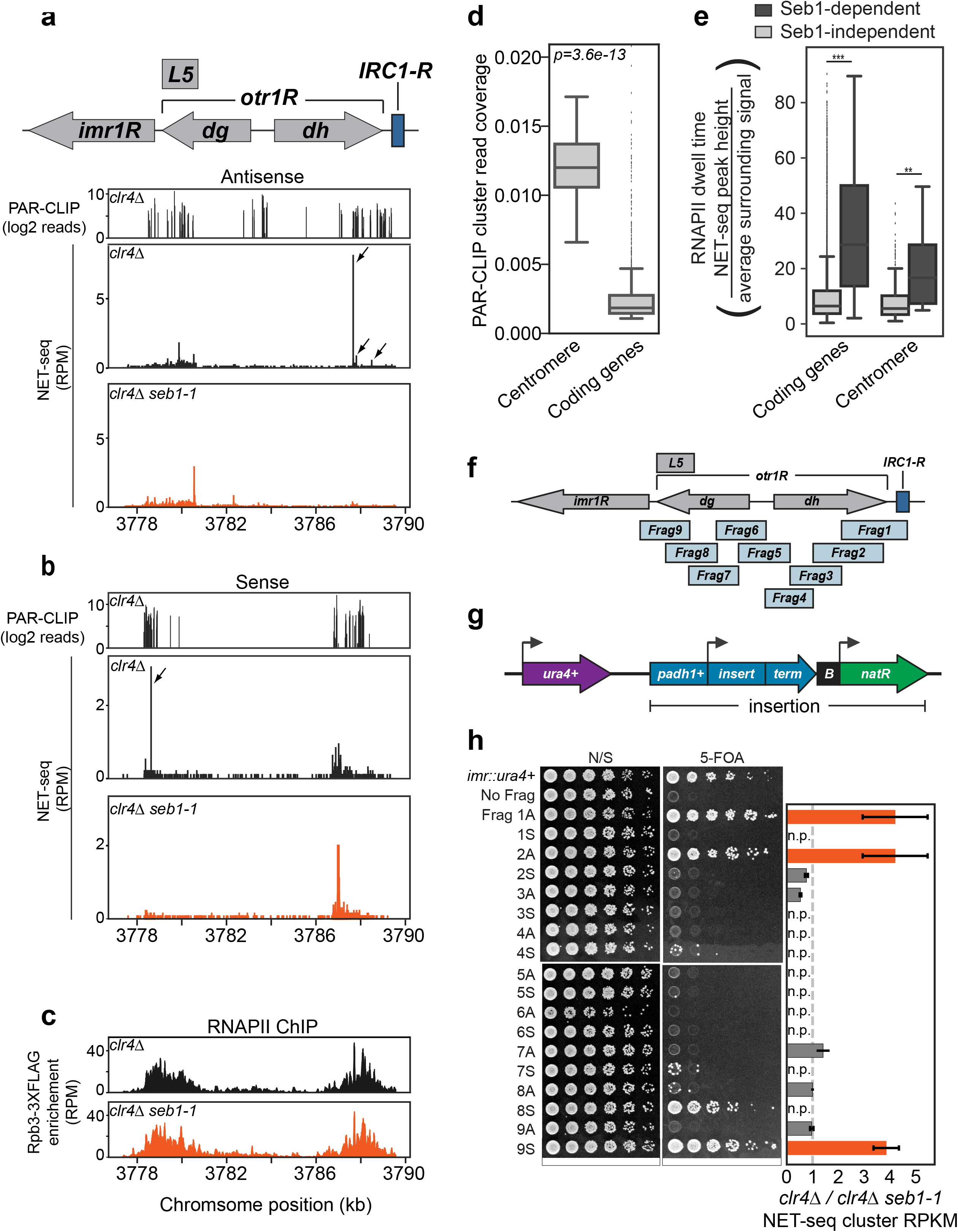
Seb1 directly binds to pericentromeric transcripts and induces RNAPII pausing in silencing-promoting segments. **a**-**b**, Seb1 PAR-CLIP read clusters (log2 reads) and NET-seq peaks (RPM) aligned to the right arm of centromere 1 in *clr4*∆ (PAR-CLIP) or comparing *clr4*∆ (black) and *clr4*∆ *seb1-1* (red) strains (NET-seq). **a**, Reads clusters/peaks aligning to the antisense transcript. **b**, Read clusters/peaks aligning to the sense transcript. Arrows indicate locations of Seb1-dependent NET-seq peak clusters identified computationally (see Methods). **c**, ChIP-seq data of Rpb3-3xFLAG for the right arm of centromere 1. Comparing RNAPII enrichment in *clr4*∆ (black) and *clr4*∆ *seb1-1* (red). **d**, Seb1 PAR-CLIP read cluster coverage at the centromeres compared to coding genes. **e**, RNAPII dwell time analysis for centromeres and coding genes comparing Seb1-dependent (dark grey) and Seb1-independent (light grey) pauses (**p=0.0068; ***p=3.6e-59) **f**, Illustration of the right arm of centromere 1 and nine overlapping fragments analyzed **g**, Illustration of the reporter construct utilized to determine the silencing capacity of centromere fragments (**f**). *adh1*^+^ promoter (*padh1*^+^); a bidirectional terminator (*term*); B-boxes boundary element (*B*); nourseothricin-resistance gene (*natR*). Construct was placed downstream of *ura4*^+^. **h**, Silencing assays for each fragment in antisense (A) or sense (S) orientations (spotting assay). Cells were plated on non-selective YS medium (N/S) and YS medium supplemented with 5-FOA (5-FOA). Controls for plating were a strain encoding a single functional *ura4*^+^ gene placed in the innermost repeats of centromere 1 (*imr::ura4*^+^), and a strain with the construct in (**g**) containing no fragment (No Frag). Ratios of *clr4*∆/*clr4*∆ *seb1-1* for NET-seq clusters in each fragment in antisense and sense transcription units (bar graph). Value of 1 represents no change in NET-seq signal (dotted line); a value >1 represents NET-seq clusters in *clr4*∆ that are reduced in *clr4*∆ *seb1-1*. Silencing-competent fragments are orange; silencing-deficient fragments are in dark grey; n.p. denotes no peaks.

**Figure 3.**
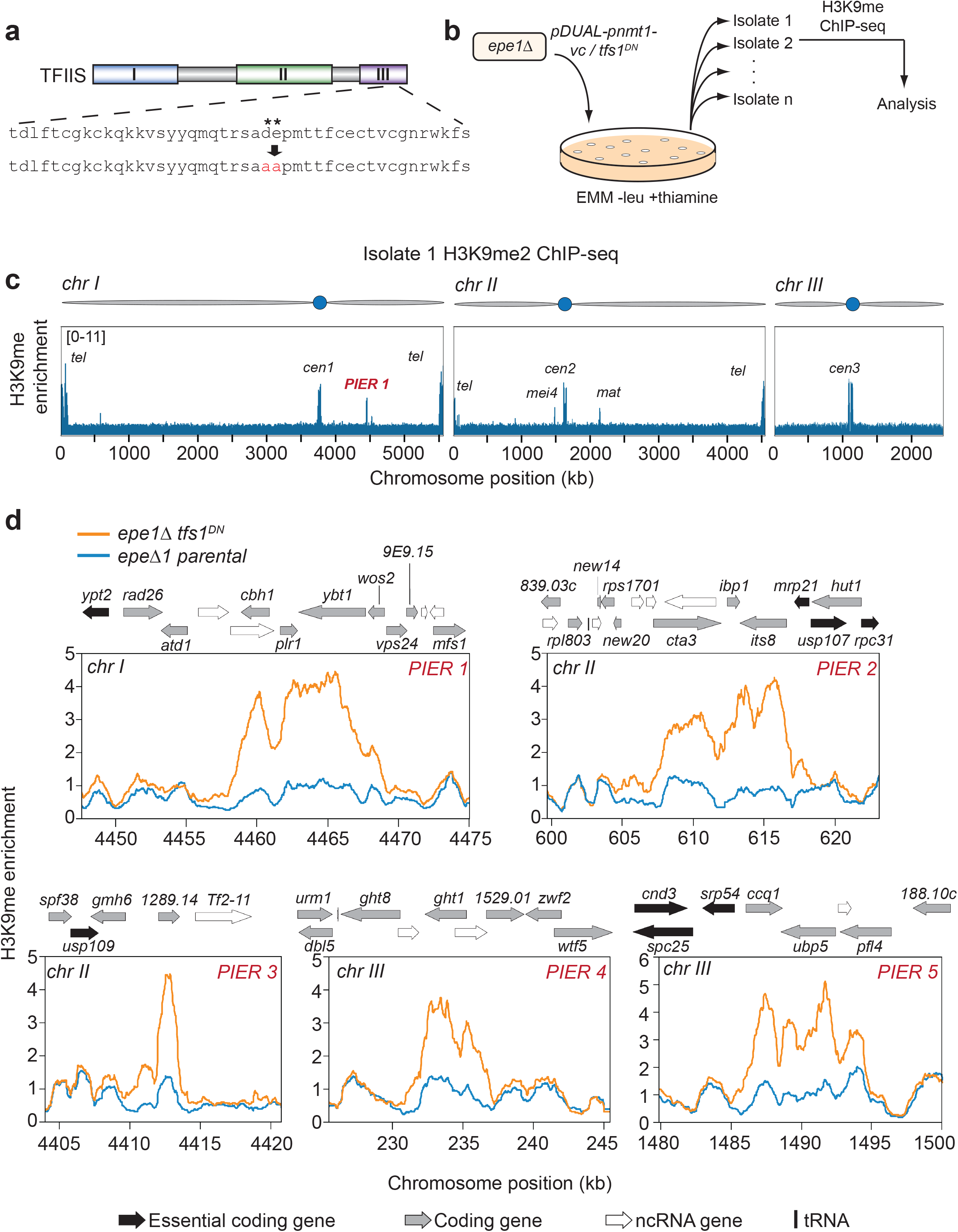
TFIIS^DN^ induces ectopic heterochromatin formation. **a**, Representation of the domains present in TFIIS. Two acidic residues in domain III (denoted by *) were mutated to alanine residues to produce TFIIS^DN^. **b**, Pipeline for isolating TFIIS^DN^ expressing cells and analysis of genome wide H3K9me2. *S. pombe epe1∆* strains were transformed with *tfs1*^*DN*^ controlled by *nmt1*^+^ thiamine repressible promoter (*pDUAL-*pnmt1^*+*^-tfs1^*DN*^) or a vector control (*pDUAL-pnmt1^+^-vc*) and selected on EMM –leu +thiamine plates. 13 isolates for each *epe1*∆ tfs1^*DN*^ and 15 for *epe1*∆ vector control strains were collected. ChIP-seq for H3K9me2 was conducted and analyzed for each isolate as well as a parental strain for each set of isolates. **c**, Genome wide representation of H3K9me2 ChIP-seq enrichment for *epe1*∆ tfs1^*DN*^ Isolate 1. PIER = <u>P</u>ause-<u>I</u>nduced <u>E</u>ctopic heterochromatic <u>R</u>egion. **d**, Genome browser images of PIERs 1 through 5 that were observed from *epe1*∆ tfs1^*DN*^ isolates. Each plot contains the H3K9me2 enrichment of the parental *epe1*∆ strain (blue) and the *epe1*∆ tfs1^*DN*^ strain (orange). Genome features are displayed above each browser image. Essential coding gene = black arrow; coding gene = grey arrow; ncRNA = white arrow; tRNA = black bar.

To compare the binding of Seb1 across transcript classes, we computed the fraction of RNA covered by Seb1 PAR-CLIP read clusters. We observed a ~12-fold higher PAR-CLIP cluster coverage for pericentromeric repeat intervals than for coding gene intervals (Figure 2d and Extended Data Figure 4a). Non-coding RNAs display the highest coverage at a mean level ~100-fold higher than that of coding-genes (Extended Data Figure 4b).

**Figure 4.**
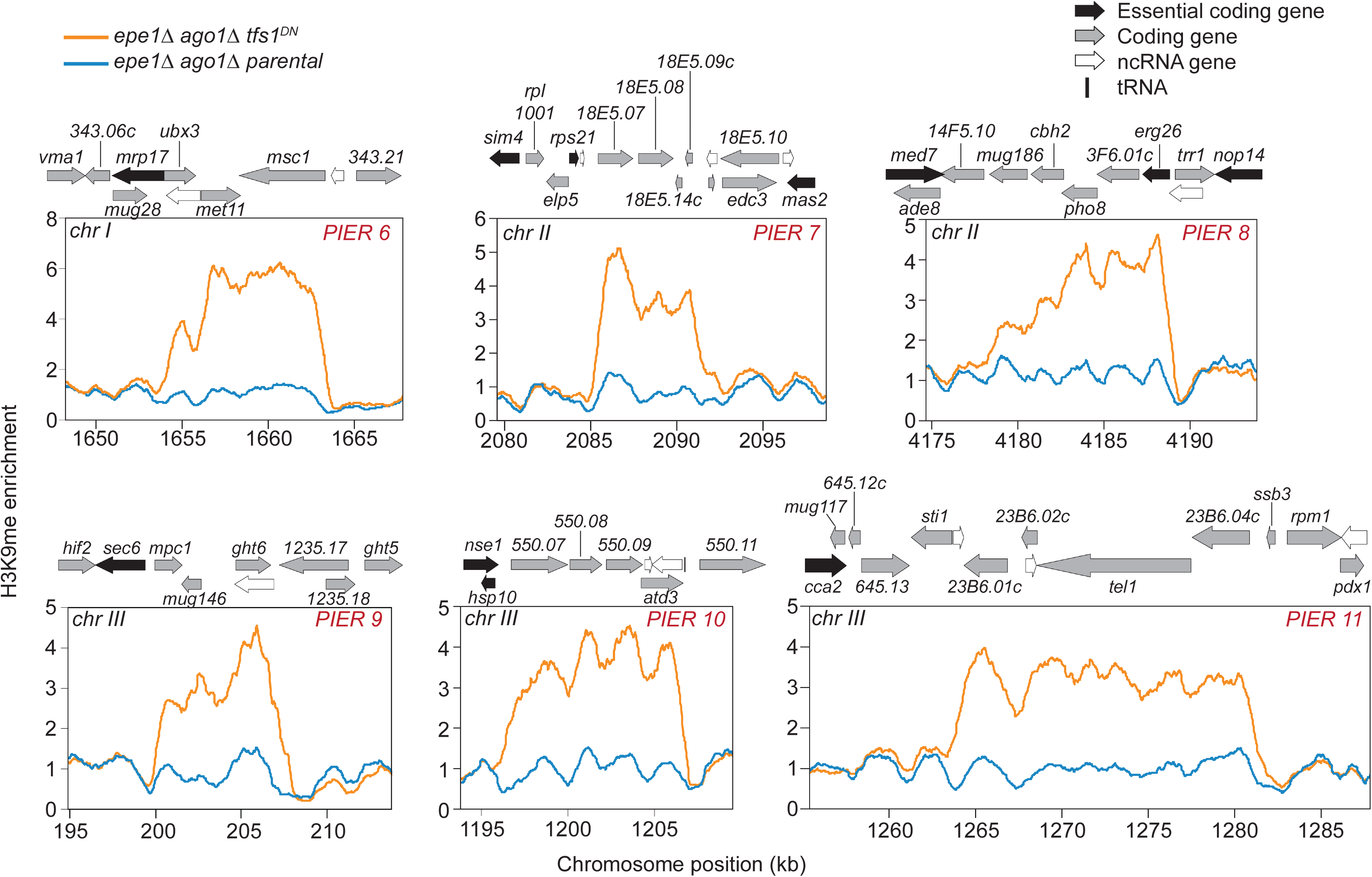
TFIIS^DN^-induce ectopic heterochromatin is RNAi-independent. Genome browser images of PIERs 6 through 11 that were observed from *epe1*∆ *ago1*∆ *tfs1*^*DN*^ isolates. *epe1∆ ago1∆* strains were transformed with *pDUAL-pnmt1*^+^-*tfs1*^*DN*^ and H3K9me2 ChIP-seq was conducted. Genome-wide H3K9me2 enrichment was analyzed and 6 PIERs (PIER 6 though 11) were observed from *epe1*∆ *ago1*∆ *tfs1*^*DN*^. Each plot contains the H3K9me2 enrichment of the parental *epe1∆ ago1∆* strain (blue) and the *epe1*∆ *ago1*∆ *tfs1*^*DN*^ strain (orange). Genome features are displayed above each browser image. Essential coding gene = black arrow; coding gene = grey arrow; ncRNA = white arrow; tRNA = black bar.

We next examined the NET-Seq profiles of pericentromeric heterochromatin sequences of *clr4*∆ and *clr4*∆ *seb1-1* strains (replicate experiments were conducted). Pericentromeric regions harbor detectable NET-Seq signal in *clr4*∆ cells despite a low level of polymerase engagement at any single nucleotide (Figure 2a and b – middle panels). The signal overlaps with regions of high Seb1 PAR-CLIP cluster coverage (Figure 2a and b – top panels). Notably, a handful of discrete peaks indicative of pausing were observed, and the largest were lost in the *seb1-1* mutant for both antisense and sense signals (Figure 2a and b – bottom panels and Extended Data Table 2). RNAPII densities at the pericentromeric regions are comparable between *clr4*∆ and *clr4*∆ *seb1-1* strains as assessed by ChIP-seq analysis of Rpb3-3xFLAG (Figure 2c and Extended Data Figure 4c), indicating that decreases in NET-seq peak intensities caused by the *seb1-1* mutation are not trivially explained by loss RNAPII recruitment (*n.b.* rare pauses may not impact overall PolII density as measured by ChIP-seq). Notably, calculation of polymerase dwell times^23^ at centromeres and across the genome and genotypes revealed that Seb1-dependent pauses are significantly longer-lived on average than Seb1-independent pauses (Figure 2e and Extended Data Figure 4d). These data reveal detectable Seb1-dependent RNAPII pauses in pericentromeric sequences.

Previous studies identified two segments of pericentromeric DNA that can trigger heterochromatin (*L5*^24,25^ and *Frag1*^15^ respectively). *Frag1* defined a segment that requires both RNAi and Seb1 for its activity^15^. To compare activity of pericentromeric fragments to their transcription properties described above, we extended this analysis using a system we employed previously^15^. The *cen1R* region was divided into nine overlapping fragments (Figure 2f and Extended Data Table 3). Each fragment was placed downstream of an *adh1*+ promoter (*padh1*^+^) in either forward or reverse orientation, and upstream of a transcription terminator. This insert was then placed downstream of *ura4*^*+*^ (Figure 2g). Silencing of *ura4*^*+*^ was determined using YS-FOA plates, which selects for *ura4+* repression. The insert of Fragment 1 (*Frag1*) displays silencing activity; this construct was used previously to isolate the *seb1-1* mutant and was shown to require the *padh1*^*+*^ promoter for silencing activity^15^. Three additional fragments exhibit strong silencing activity and each is functional in only one orientation (Figure 2h; *Frag2A, 8S* and *9S*). Thus, these pericentromeric regions harbor a transcription-dependent, orientation-specific signal capable of triggering silencing. To examine the relationship of these regions to those that display detectable Seb1-dependent pauses, we identified clusters of NET-seq peaks (see Methods) and computed the total read density of these clusters within each fragment. A comparison of *clr4*∆ to *clr4*∆ *seb1-1* strains revealed significant correlation (χ^2^=12.6, p<0.001) (Figure 2h – *Frag1A*, *Frag2A*, and *Frag9S*; Extended Data Figure 5a-c). The exception, *Frag8S,* displays silencing activity but no detectable Seb1-dependent NET-seq peak clusters (although it does display Seb1-dependent NET-seq signal; Figure 2h and Extended Data Figure 5d). The heterochromatin assembly activity of this fragment may be pause-independent or, the relevant RNAPII pauses may be below the sensitivity of NET-seq. These data indicate that Seb1 directly recognizes *dg* and *dh* RNAs and induces detectable pausing in centromere fragments that display silencing activity.

The shared defect of the viable *seb1-1* allele in both heterochromatin assembly and RNAPII pausing suggests that pausing signals the assembly of heterochromatin. However, given that Seb1 may have other activities impacted by *seb1-1*, we sought an orthogonal test of the role of pausing. Thus, we pursued an alternative strategy of testing whether increasing RNAPII pausing *per se* could be sufficient to trigger heterochromatin assembly and if so whether such an activity required RNAi. We exploited the conserved elongation factor TFIIS which binds paused RNAPII complexes and stimulates RNA hydrolysis by RNAPII, enabling restart^26^. It is thought that all genes are subject to this type of rescue mechanism as pausing is a ubiquitous feature of transcription. Two conserved acid residues in domain III of TFIIS are required for catalysis^27^. Mutation of these residues to alanine prevents the cleavage of the RNA by RNAPII^27^, ultimately resulting in polymerase trapped in a lethal paused/backtracked state^28,29^. We introduced the corresponding D274A and E275A mutations in the TFIIS gene *tfs1*+, creating a dominant negative allele *tfs1*^*DN*^ (Figure 3a)^30^. To overcome lethality of this allele^28,29^, we placed it under control of an *nmt1*^*+*^ thiamine-repressible promoter and inserted it at the *leu1*^*+*^ locus^31^, enabling concerted expression of *tfs1*^+^ and tfs1^*DN*^. We conducted replicate NET-seq analysis on *clr4*∆ and *clr4*∆ tfs1^*DN*^ cells under inducing conditions. Genome-wide analysis of 5’ and 3’ traveling ratios indicated TFIIS^DN^-dependent increases in RNAPII pausing at gene 5’ ends and a more modest effect at 3’ ends (Extended Data Figure 6a-b).

To test whether stabilizing endogenous RNAPII pauses in this manner triggers *de novo* ectopic heterochromatin, we performed H3K9me2 ChIP-seq analysis on *tfs1*^*DN*^ strains; however, we observed no effects in this background (data not shown). Heterochromatin components are limiting and antagonized by anti-silencing factors, particularly Epe1, which actively removes the H3K9me mark^32–34^. Thus, we constructed *epe1∆* tfs1^*DN*^ mutant strains or *epe1∆* strains carrying an integrated vector-only control (*epe1∆-vc*), collected multiple strain isolates for each, and performed H3K9me2 ChIP-seq on all isolates (Figure 3b). Because high-level TFIIS^DN^ expression is lethal in *epe1*∆ cells (Extended Data Figure 6c – middle panel), experiments were performed in the presence of thiamine, enabling viability (Extended Data Figure 6c – right panel). Consistent with a slight fitness defect under these conditions (Extended Data Figure 6c – right panel), RNA-seq analysis revealed that 13% of the transcript pool from the *tfs1* genes arise from the *tfs1*^*DN*^ allele and 87% arise from the wild-type allele when cells are grown in thiamine (Extended Data Figure 6d), indicating leaky repression. The ChIP-seq data obtained from strain isolates were examined for H3K9me peaks (see Methods). We filtered the results for well-established heterochromatic recruitment sites (including HOODs, Islands, meiotic genes, Epe1-bound genes etc; see Methods), as these genomic regions have an intrinsic propensity (e.g. via RNAi or the RNA elimination machinery) to nucleate H3K9me.^35–37^

Remarkably, of 13 isolates derived from *epe1*∆ *tfs1*^*DN*^ parents analyzed by H3K9me ChIP-seq, five separate isolates harbor a distinct ectopic region of heterochromatin, which we termed <u>P</u>ause-<u>I</u>nduced <u>E</u>ctopic heterochromatic <u>R</u>egion (PIER) (Figure 3c and d; Extended Data Figure 7). No novel ectopic heterochromatic loci were observed by whole genome H3K9me2 ChIP-seq in the 15 *epe1*∆-*vc* strains (χ^2^=7.02, p=0.0082). PIERs range in size from ~3 to ~15 kb, and each PIER was unique. Three PIERs are bounded by an essential gene on at least one side of the locus suggesting that selection likely prevents observing PIERs that assemble over essential genes (Figure 3d PIERs 2, 3, and 5); this implies that our approach may underestimate the propensity of PIER formation. H3K9me enrichment at known heterochromatin nucleation sites is unrelated to *tfs1* genotype and summarized for all strains in **Extended Data Figure 8**. These results indicate that TFIIS^DN^-induced RNAPII pausing can be sufficient to nucleate heterochromatin at novel sites.

To determine if H3K9me at PIERs lead to repression, we conducted RNA-seq analysis on *epe1*∆ *tfs1*^*DN*^ isolate 2 (containing PIER 2) (Extended Data Figure 9a and Extended Data Table 4). We observed a significant decrease (p<0.01) in two of the three genes present in PIER2, *cta3*^+^ and *its8*^+^ (Extended Data Figure 9a and Extended Data Table 4). TFIIS^DN^ expression does not alter the expression of known heterochromatin factors (p<0.01, |log_2_(fold-change)|>1; Extended Data Figure 9b; Extended Data Table 4).

Four out of five PIERs contain overlapping convergent genes that have the potential to form double stranded RNA (Figure 3d; PIERs 1, 2, 4, and 5). As double-stranded RNA can trigger heterochromatin in *cis* via the RNAi pathway^38^, we determined whether the *de novo* establishment of PIERs requires RNAi. We constructed *epe1∆ ago1∆* strains by disrupting *ago1*^+^ in the *epe1*∆ mutant. Initial ChIP-seq analysis of three isolates revealed two that display ectopic heterochromatin at the *clr4*^+^ locus and reduced levels of H3K9me at constitutive heterochromatic loci (Extended Figure 10 and Extended Data Figure 11 b and c top panels). Such adaptive silencing of *clr4*^+^ has been described previously in *epe1∆ mst2∆* strains^37^ and evidently can occur in strains lacking Epe1 and RNAi. Remarkably, upon integration of *tfs1*^*DN*^ into these two *epe1∆ ago1∆* strains, H3K9me2 ChIP-seq revealed establishment of heterochromatin at constitutively silenced loci (e.g. centromeres) and its loss at the *clr4*^+^ locus (Extended Data Figure 11b and c – isolates 1, 3, 4, and 5). This observation indicates that pericentromeric repeats harbor the ability to respond to RNAPII pausing and assemble heterochromatin independently of RNAi. Accompanying these changes, six out of the 12 *epe1*∆ *ago1*∆ *tfs1*^*DN*^ isolates subjected to whole-genome analysis acquired PIERs (Figure 4; Extended Data Figure 11). Five of six of these PIERs are bounded by an essential gene on at least one side of the region similar to that seen in the *epe1*∆ *tfs1*^*DN*^ (Figure 4; PIERs 6-10), consistent with the notion that essential genes limit our ability to observe ectopic heterochromatin. Again, each PIER was unique. Thus, PIERs can be triggered by RNAPII pausing even in cells lacking a functional RITS complex.

Our results indicate that Seb1, a conserved RNAPII-associated RNA binding protein that mediates RNAi-independent heterochromatin assembly in *S. pombe*^15^, is enriched on pericentromeric ncRNA transcripts relative to coding sequences and promotes long-lived RNAPII pauses. Remarkably, pausing is sufficient to trigger ectopic heterochromatin assembly in an RNAi-independent fashion, indicating that this is a relevant activity of Seb1 in promoting heterochromatin assembly. Binding of Seb1 to euchromatic ncRNAs (e.g. snRNAs) is not associated with detectable heterochromatin assembly; this may be due to high levels of transcription, which induces anti-silencing histone marks and histone turnover, both of which antagonize silencing^1^. Alternatively, the repetitiveness of pericentromeric sequences may also contribute to specificity by producing a threshold density of paused polymerases within a discrete genomic interval. Testing these and other possibilities will require the development of synthetic biology tools that enable the programming of pauses of defined length at defined sites and at define levels of transcription, a task beyond current technology. Our data are germane to the observation that mutations in the Paf1 complex (Paf1-C), a multifunctional elongation complex that binds cooperatively to RNAPII with TFIIS^39^, enables synthetic hairpin RNAs to trigger heterochromatin in *trans* and increases heterochromatin spreading in *S. pombe*^40–42^. While the elongation-promoting activity of Paf-C has been suggested to limit heterochromatin by limiting targeting of RITS to the nascent transcript^43^, it may also act via RNAi-independent mechanisms as we find that increased RNAPII pausing can trigger H3K9me independently of RNAi. Seb1-triggered RNAPII pausing may drive heterochromatin assembly by promoting the heterochromatic stalling of replisomes associated with CLR-C through RNAPII-replisome collisions as proposed^44,45^ (Extended Data Figure 12). Analogous concepts have been put forth in *S. cerevisiae* where tight protein-DNA interactions are sufficient to trigger recruitment of the SIR complex^46^. Consistent with this hypothesis, such transcription-replication conflicts are limited by Paf1-C^47^ which inhibits heterochromatin assembly, while slowing of replisome progression enhances heterochromatin spread^48,49^. It has also been proposed that the 5’→3’ RNA exonuclease Dhp1 (related to *S. cerevisiae* Rat1/Xrn2), which is required for RNAi-independent heterochromatin assembly, recruits the silencing machinery via a physical interaction with CLR-C^50,51^. Because RNAPII pausing enhances recruitment of Xrn2^52,53^, and Seb1 copurifies with the Dhp1^14,19^, Seb1-induced pausing may promote heterochromatin assembly via this mechanism as well (Extended Data Figure 12). Our model also readily accommodates genetic observations that null mutants in *S. pombe* RNAPII elongation factors suppress the H3K9me defect of RNAi mutants^16,41^, as well as analogous observations for mutants in RNA biogenesis factors^16^ as these factors also promote transcriptional elongation^54^. Weak cleavage-polyadenylation signals promote heterochromatin assembly^55^, which is predicted to result in accumulation of paused RNAPII at *S. pombe* Downstream Pause Elements^56^. Another key factor recruited to pericentromeric regions by Seb1 is remodeling/HDAC complex SHREC^57^. Thus, Seb1-paused RNAPII may promote heterochromatin assembly through multiple mechanisms. Other fungi, such as *Neurospora crassa* and *Cryptococcus neoformans,* as well as somatic mammalian cells, do not require RNAi for heterochromatin assembly^58–60^; thus, the RNAi-independent mechanisms discussed here may apply to such systems. Given the tight coupling of this heterochromatin signal to RNAPII activity, it is tempting to speculate that Seb1-mediated pausing may have evolved from a surveillance mechanism for silencing foreign DNA. Finally, pathogenic triplet repeat expansions in the Friedreich Ataxia gene *FXN* concomitantly display a block to transcriptional elongation and the appearance of H3K9me on *FXN*^61,62^, raising the possibility that pause-induced heterochromatin underlies disease pathogenesis.

## Methods

### Yeast strains, plasmids, and media

A list of all *S. pombe* strains and plasmids used in this study is provided in Extended Data Table 5. Cells were grown at 30°C in synthetic complete medium (SC) with adenine and amino acid supplements with reduced levels of uracil (150mg/L) for PAR-CLIP, or in Edinburgh minimal medium (EMM) supplemented with adenine, uracil, and the appropriate amino acids with or without thiamine (15µM) for NET-seq and ChIP-seq.

### NET-seq

NET-seq experiments were conducted as previously described^63^ with minor alterations for *S. pombe. S. pombe* cultures were grown in 1L EMM without thiamine to an OD600 of 0.7 and harvested via filtration and flash frozen in liquid nitrogen. Lysis and immunoprecipitation was conducted as previously described^64^. Adaptor ligation was performed using random hexamer-barcoded adaptors. All strains were analyzed in duplicate and sequencing was conducted on a HiSeq 4000 platform.

### RNA-seq

Strains were grown in YS media + 3% Glucose overnight to OD_600_ = 0.7. Cells were harvested by centrifugation, washed twice with ice-cold water and flash-frozen. Pellets were resuspended in 1ml Trizol (Thermo Fisher Scientific, #15596026). 0.5 mm zirconia-silica beads (BioSpec, #11079105z) were added and lysis was accomplished by three cycles of bead beating for 90 seconds on high (Bead Ruptor 12 Homogenizer, OMNI International). Following centrifugation at 14,000 rpm for 10 min at 4°C, the supernatant was transferred to a microcentrifuge tube, extracted once with chloroform, and precipitated with isopropanol. Following resuspension and re-precipitation with isopropanol, pellets were washed with 75% ethanol and air dried for 30 minutes. Pellets were resuspended in 150µl RNAse-free water.

1mg of total RNA was used to isolate mRNA using PolyATtract^®^ Systems III and IV (Promega, #Z5310) according to the manufacturer’s instructions. Input RNA quality and mRNA purity were verified by Bioanalyzer RNA 6000 Pico kit (Agilent, #5067-1513). To address the issue of genomic DNA contamination in RNA samples, we use Zymo RNA Clean & Concentrator Kit 5^®^ (Zymo Research, #11-326) according to the manufacturer’s instruction. A sequencing library was constructed using NEBNext Ultra Directional RNA library Prep Kit for Illumina (New England Biolab, #E740S). Librairies were analyzed for quality and average size on Bioanalyzer High Sensitivity DNA kit (Agilent, #5067-4626). The sequencing was performed on an Illumina HiSeq 4000 platform.

### PAR-CLIP

PAR-CLIP experiments were conducted as a combination of PAR-CLIP and CRAC protocols^65,66^ in two replicates. Cells were grown in 2L of SC medium to an OD_600_ of 0.75. 4-thiouracil (Sigma, #440736-1G) was added to a final concentration of 1.3mM for 15 minutes in the dark at 30°C. Samples were immediately crosslinked using the UV Power-Shot Handheld UV Curing System (SPDI UV) at a 365 nm wavelength for 15 minutes while continuously stirring. Samples were collected by filtration, resuspended in 6mL of buffer TMN150 (50mM Tris-HCl pH 7.8, 150mM NaCl, 1.5 mM MgCl2, 0.5% NP-40, 5mM beta-mercaptoethanol) and frozen a “yeast popcorn” by dropwise addition to liquid nitrogen. This material was lysed by ball mill (15 Hz, 3 minutes, 5 cycles) (Mixer Mill MM 301, Retsch). CRAC was then preformed from this point on as previously described^65^ with the exception of gel extraction, which was conducted by electroelution of the gel piece containing the radioactively labeled RNA sample using D-TubeTM Dialyzer Midi tubes (EMD Millipore, #71507-3). Electroelution was carried out in 1X MOPS SDS PAGE Buffer (5mM MOPS, 50mM Tris, 0.1% SDS, 1mM EDTA) and run for 2hrs at 100V. The isolated sample was transferred to a fresh tube and 100ug of Proteinase K (Sigma, #P2308) was added to the sample. Procedures post-Proteinase K treatment were conducted as previously described^65^ and samples were sequenced on the HiSeq 4000 platform.

### Spotting Assay

Strains were grown overnight to saturation and diluted to OD_600_ of 1. Serial dilutions were performed with a dilution factor of 5. For *ura4*^+^ silencing assays, cells were grown on non-selective and 5-fluoroortic acid (FOA) (2 g/L) YS plates at 30°C for 3 days. For *tfs1*^*DN*^ induction assays, cells were grown on YS, EMM -leu -thiamine, or EMM -leu +thiamine, for 3 days.

### ChIP-seq

ChIP-seq was conducted as previously described^67^. ChIP-Seq immunoprecipitations were performed with 10 µg anti-H3K9me2 (Abcam, #ab1220), and 10 µg anti-FLAG (Sigma, #P3165). Samples were sequenced on a HiSeq 4000 platform.

### NET-seq analysis: Genome alignment

For NET-seq analysis, adapter sequences were removed and reads were flattened to remove sequence duplicates. Barcoded reads were then mapped to the *S. pombe* genome^68^ using BOWTIE^69^ to align and omit any sequence reads that were misprimed during the reverse transcription step of NET-seq and thus lack a barcode using the following flags: -M1 --best --strata. Unaligned files were collected for further analysis. Barcodes were removed, and the new unique, debarcoded reads were realigned to the genome using the following flags in BOWTIE: -M1 --best --strata.

### NET-seq analysis: Cluster finding

High-density regions of NET-seq signal were defined across centromeres and coding regions to compare NET-seq density between genotypes. First, NET-seq peaks were discovered by calculating robust Z-scores (based on median and median absolute deviation) from the log2 transform of the number of reads starting at each position in the defined region (centromere fragment or transcript). Positions with a robust Z-score of at least 2 and at least 10 unique reads were considered peaks. Next, peaks were clustered together using a sliding window (width=50, increment=10). The density of the cluster is calculated as the number of reads in the cluster divided by the size of the cluster in kilobases (kb).

To determine cluster densities for each fragment derived from the right arm of centromere 1, the sum of cluster densities was normalized to the sum of all densities in each sample. Error bars represent the range of two replicates.

### NET-seq analysis: Traveling ratio

Traveling ratios were calculated for every non-overlapping annotated transcript at least 1000 nt in length according to the Pombase annotation^68^. Transcripts with fewer than 50 total reads were excluded from the analysis. The traveling ratio was determined for a 0.5 kb window either immediately after the transcription start site (5’ traveling ratio) or immediately before the cleavage polyadenylation site (3’ traveling ratio). Transcripts <1kb in length were omitted from this analysis to ensure the 0.5kb 5’ and 3’ windows used for each traveling ratio do not overlap. Reads were counted in this window and across the entire transcript and then divided by the size of the window or transcript, respectively. Transcripts were clustered using K-means (sklearn.cluster.Kmeans, 3 centroids) based on the travelling ratio at each end of the *clr4*∆ and *clr4*∆ *seb1-1* mutant on one replicate. P-values for each cluster for the difference between the *clr4*∆ and *clr4*∆ *seb1-1* traveling ratio distributions were determined by KS test for each pair of replicates. Traveling ratio CDF plots were similar between replicates and a single replicate is presented.

### NET-seq analysis: Dwell time

Dwell time was determined by normalizing peak height to the average NET-seq signal density of the surrounding 100 nt. NET-seq peaks with at least a two-fold decrease from *clr4*∆ to *clr4*∆ *seb1-1* in both replicates were considered Seb1-dependent. P-values were determined by KS test.

### RNA-seq analysis

Analysis was performed using TopHat^70^ and DESeq2^71^. Changes in transcript expression levels required >2-fold change in mutants compared to wildtype to be considered significantly changed enough to have a functional consequence. Data analysis was performed on 2 replicates per condition.

To determine the fraction of reads derived from the expression of *tfs1*^*DN*^ we divided the total number of reads that specifically aligned to the mutated region of the *tfs1*^*DN*^ allele by the total number of reads (both WT and mutant alleles) that aligned to this same region.

### Seb1 PAR-CLIP data analysis by PARalyzer

For PAR-CLIP analysis, adapter sequences were removed and reads were mapped to the *S. pombe* genome^68^ using BOWTIE^69^, allowing for three mismatches with the following flags: -M1 -v3 --best --strata. Seb1 binding site read clusters were identified with PARalyzer^21^. Reads of < 20nt were omitted, and read clusters required at least 10 reads and at least two T→C conversions per cluster to be called as a Seb1 binding site. The PARalyzer OUTPUTCLUSTERSFILE file was converted to a genome browser readable file (.bam) for analysis. PAR-CLIP cluster coverage was calculated as the fraction of the interval of interest harbouring covered by a PARalyzer-called PAR-CLIP cluster (centromere arm, coding gene or ncRNA). P-values were determined by KS test.

### DREME motif analysis

DREME^22^ motif discovery for short, ungapped sequences was utilized to find Seb1-specific binding motifs. PAR-CLIP clusters – that were flattened to remove identical, recurrent sequence clusters originating from all three centromeric regions – were subjected to motif analysis. A shuffled sequence set created from the input sequences was utilized as a control.

### ChIP-seq analysis

ChIP-seq analysis was conducted as previously described^67^. Briefly, adaptor sequences from ChIP-seq sequencing libraries were removed and reads <20nt were omitted. Reads were aligned to the *S. pombe* genome^68^ using BOWTIE^69^ with the following flags: -M1 --best --strata. Aligned reads were smoothed over a 1kb window.

### ChIP-seq analysis: PIER discovery

H3K9me ChIP-seq peaks were considered as novel ectopic sites of H3K9me if two criteria were met: 1) H3K9me peaks were ≥3-fold higher than the genome background signal in the isolate, and 2) when normalized to the WCE, the H3K9me signal at the peak was ≥3-fold higher than the parental H3K9me ChIP-seq signal. A curated list of genomic regions previously observed to have a propensity to form heterochromatin in various *S. pombe* backgrounds^35–37^ was generated (Extended Data Table 6). In *epe1*∆ backgrounds, oscillation and spreading of H3K9me can occur^72^; thus, peaks within 10kb of our curated list of H3K9me nucleation sites, or within 10kb of H3K9me regions present in the parental strain, were not counted as novel H3K9me nucleation events.

### ChIP-seq analysis: H3K9me levels at HOODs, islands, meiotic genes, and PIERs

For all isolates and whole cell extracts, RPKMs for each region in Extended Data Table 6 and all PIERs were normalized to the RPKM of a 10kb window surrounding each region (5kb upstream and downstream). The ratio of H3K9me enrichment values from isolates to the whole cell extracts and plotted as a heatmap (Extended Data Figure 8)

### Data sets

All available sequencing data sets are listed in Extended Data Table 7.

## ACKNOWLEDGEMENTS

We thank the members of the Madhani lab for support and scientific discussion, Robin Allshire (The University of Edinburgh) for the gift of the *pDUAL-pnmt1*+ plasmid, and Fred Winston and Ameet Shetty (Harvard Medical School) for sharing their *S. pombe* NET-seq protocol. We thank Diana Marina, Smita Shankar, and Ken Finn for their early contributions to this work. We thank Sigurd Braun, Geeta Narlikar, Robin Allshire, Phillip Dumesic and Bassem Al-Sady for critical review of the manuscript. Research in the Madhani laboratory is supported by grants from the U.S. National Institutes of Health. H.D.M. is a Chan-Zuckerberg Biohub Investigator.

## AUTHOR INFORMATION

### Contributions

J-Y.P. and H.D.M. conceived and designed the study. J-Y.P. designed and executed ChIP-Seq, NET-Seq, and PAR-CLIP experiments. S.B. designed, executed and analysed the RNA-seq experiments. J-Y.P. performed analysis of ChIP-Seq data. J-Y.P., J.E.B. and C.H. performed computational analysis for PAR-CLIP data. J.E.B. performed computational analysis for NET-Seq data. J-Y.P. and H.D.M. wrote the manuscript.

---

**Extended Data Figure 1.**
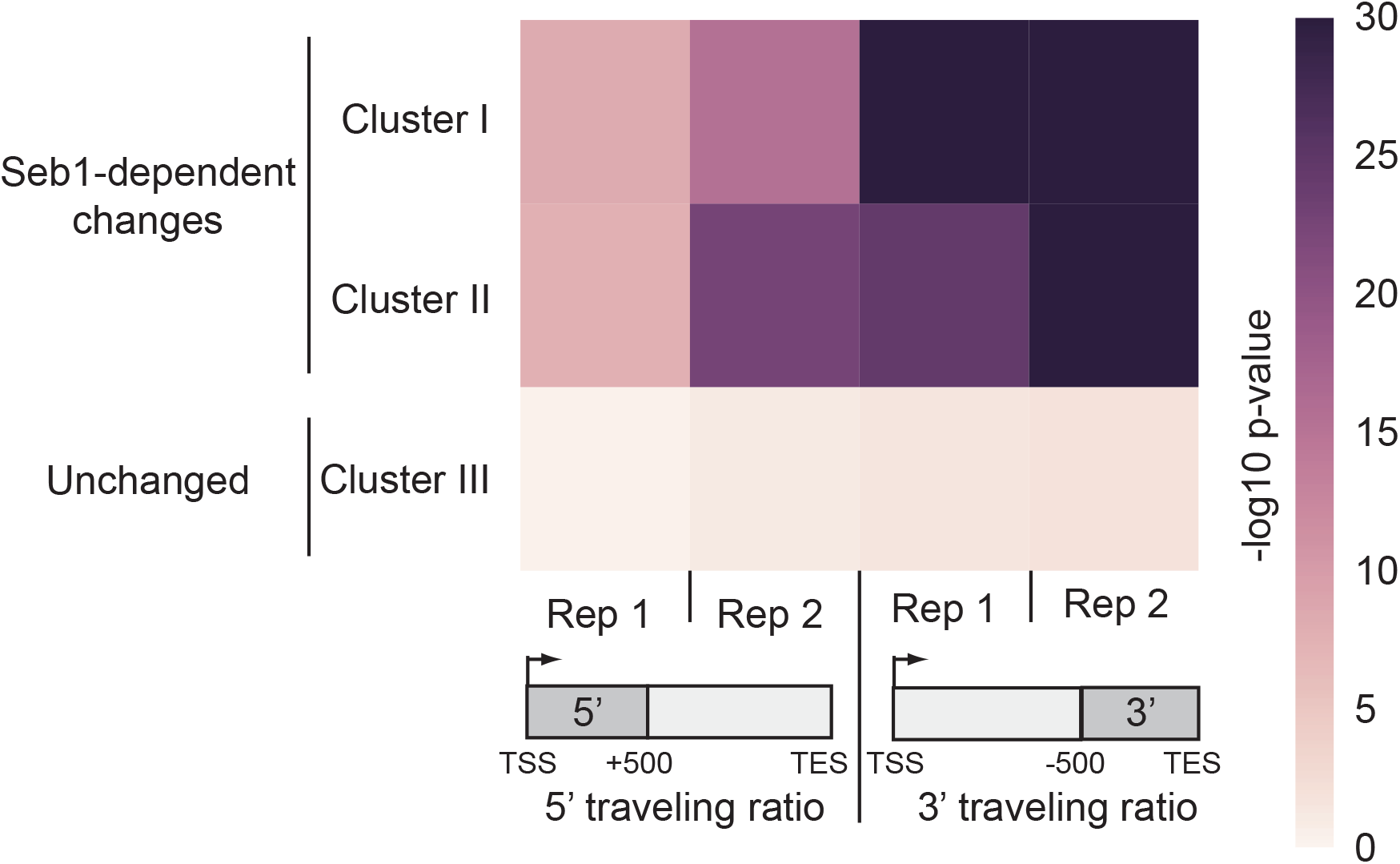
*seb1-1* mutant displays statistically significant changes in 5’ and 3’ RNAPII traveling ratios. Heat map of −log10 p-values of KS tests comparing *clr4*∆ to *clr4*∆ *seb1-1* traveling ratios from each set of clusters in (**Figure 1d**). Clusters I and II display significant Seb1-dependent changes. Each replicate is represented (Rep 1 and Rep 2) for both 5’ and 3’ traveling ratios.

**Extended Data Figure 2.**
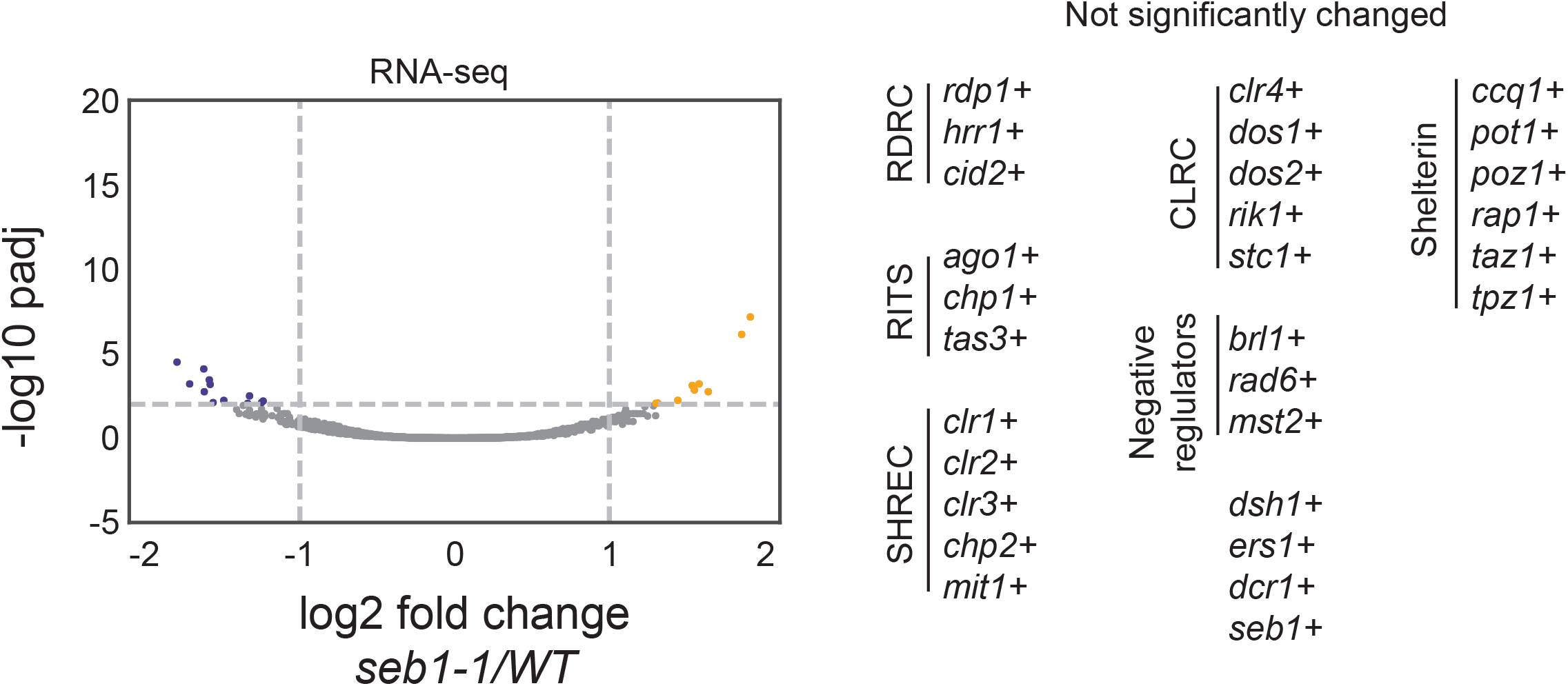
RNA-seq anlaysis of *seb1-1*. **a**, (left) RNA-seq volcano plots of log2 fold changes in transcript levels in *seb1-1* compared to WT. 15 genes have significantly lower expression in *seb1-1* (blue dots); 9 genes have significantly higher expression in *seb1-1* (orange dots). padj = adjusted p-value obtained from DESeq2. (right) No significant changes in transcript levels were observed in factors known to play a role in heterochromatin formation.

**Extended Data Figure 3.**
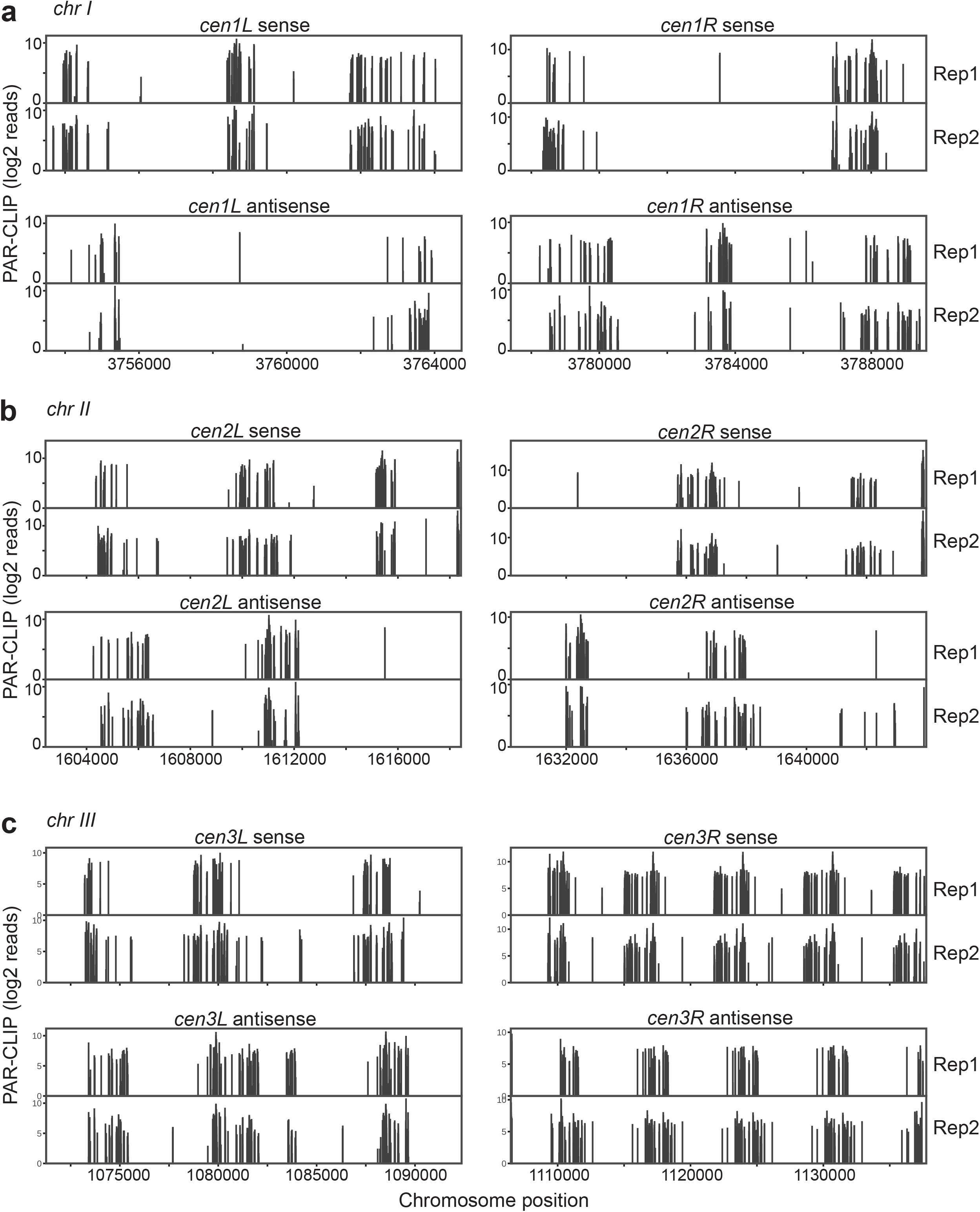
PAR-CLIP analysis of Seb1 at all three centromeres. **a**-**c**, Seb1 PAR-CLIP clusters are present at centromeres 1 (**a**), 2 (**b**) and 3 (**c**). Centromere left (cenL) and right (cenR) arms are depicted for two replicates (Rep 1 and Rep 2) in both sense and antisense orientations.

**Extended Data Figure 4.**
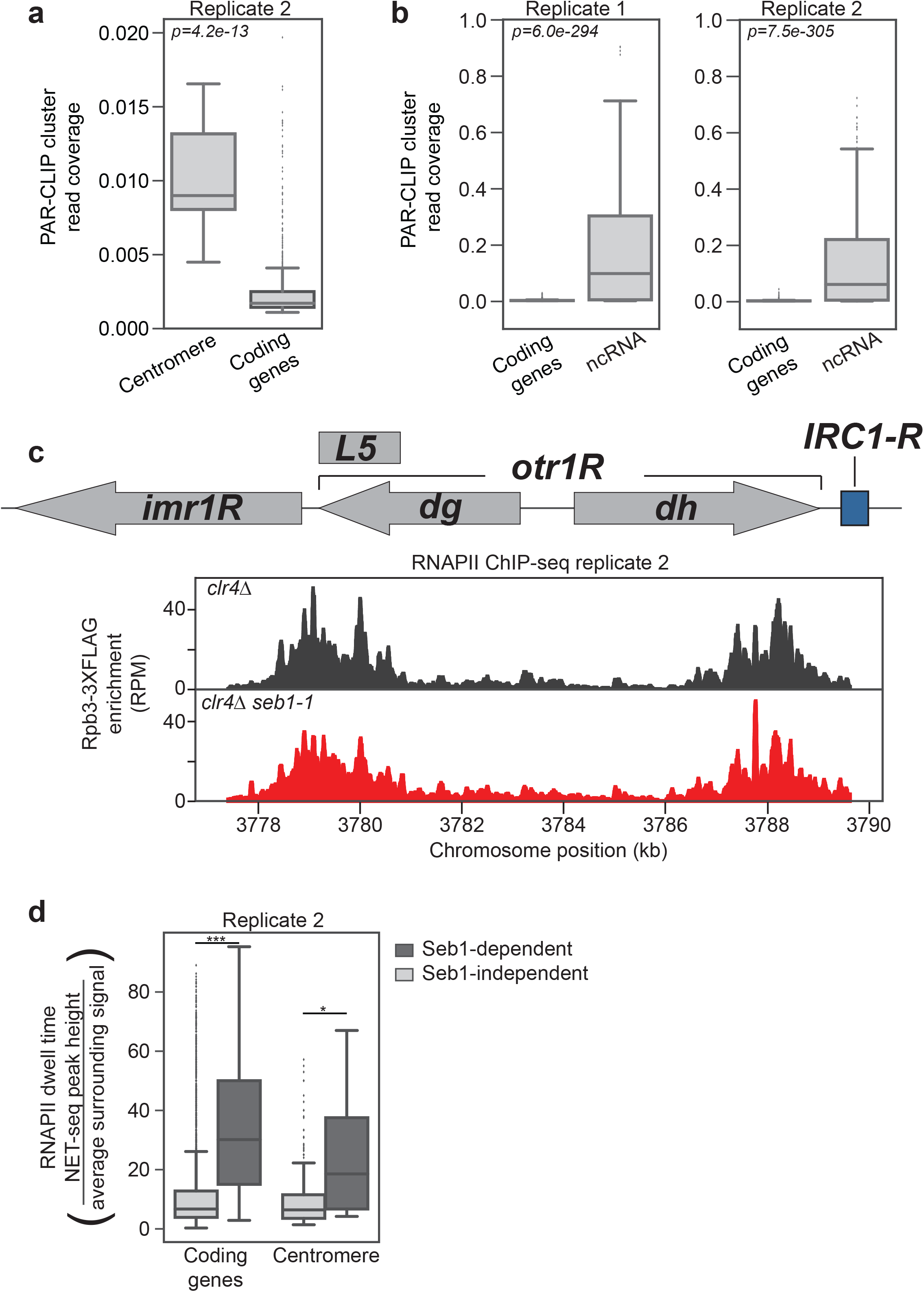
Seb1 induces RNAPII pauses with long dwell times throughout the genome. **a,** Seb1 PAR-CLIP cluster read coverage at the centromeres compared to coding genes for the second replicate. **b**, Seb1 PAR-CLIP cluster read coverage at ncRNAs (including sn/snoRNAs, tRNAs) compared to coding genes. **c**, Replicate 2 data for RNAPII levels based on ChIP-seq of Rpb3-3xFLAG with reads aligned to the right arm of centromere 1. Comparing RNAPII enrichment in *clr4*∆ (black) and *clr4*∆ *seb1-1* (red). **d**, RNAPII dwell time analysis for centromeres and coding genes in a second replicate (*p=0.026; ***p=1.8e-56).

**Extended Data Figure 5.**
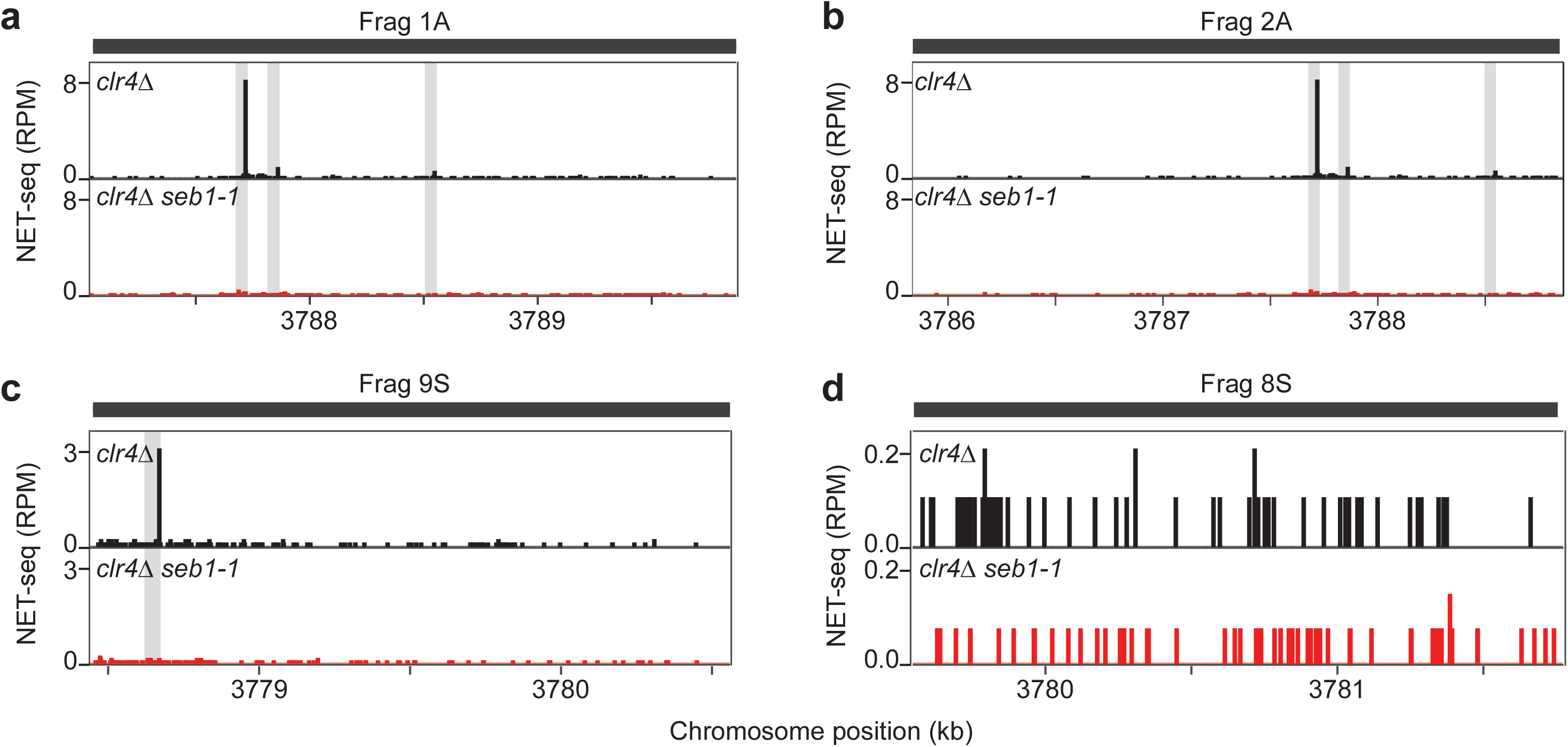
Seb1 induces pauses within centromere fragments that have a capacity to assemble heterochromatin. **a-d**, NET-seq read density (RPM) for *clr4*∆ (black) and *clr4*∆ *seb1-1* (red) aligned to the four fragments in **Figure 2h** that display a capacity to induce heterochromatin formation in the context the reporter construct shown in **Figure 2g**. **a**, *Frag1A*; **b**, *Frag2A*; **c**, *Frag9S*; **d**, *Frag8S*. Grey bars represent locations of NET-seq peaks/RNAPII pause sites present in the *clr4*∆ strain.

**Extended Data Figure 6.**
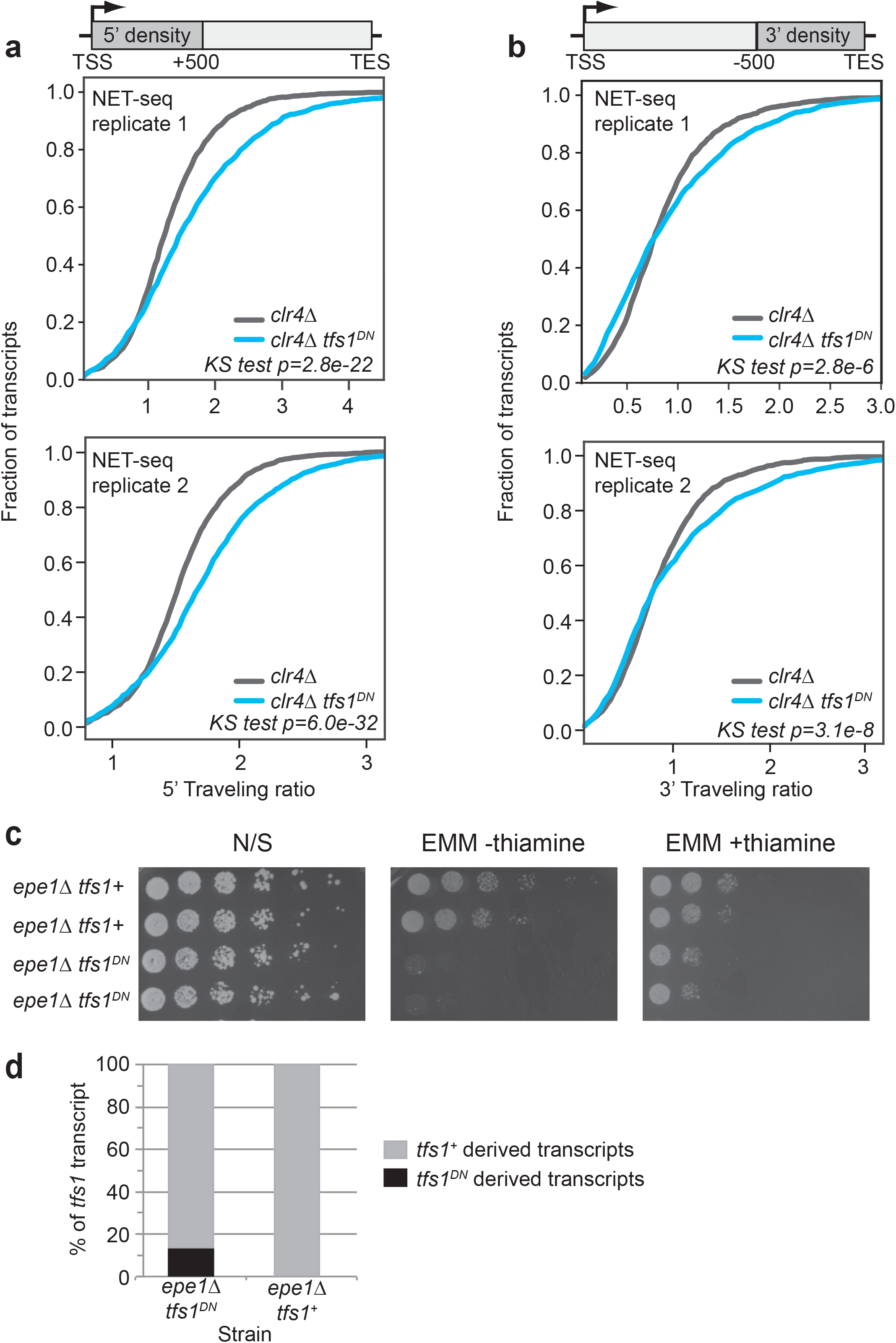
*pnmt1*^+^-*tfs1*^*DN*^ expression impacts traveling ratios genome wide. **a**, Cumulative distribution function (cdf) plots of 5’ traveling ratios for *clr4*∆ (dark grey) to *clr4*∆ *tfs1*^*DN*^ (red) NET-seq strains for two replicates. KS tests were conducted for p-values. **b**, Cumulative distribution function (cdf) plots of 3’ traveling ratios for *clr4*∆ (dark grey) to *clr4*∆ *tfs1*^*DN*^ (red) NET-seq strains for two replicates. KS tests were conducted for p-values. **c**, Growth assays on *epe1*∆ *tfs1*^+^ and *epe1*∆ *tfs1*^*DN*^ strains. Cells were plated on non-selective rich YS medium (N/S), EMM medium in the absence of thiamine (EMM -thiamine), and EMM medium containing 15uM thiamine (EMM +thiamine). **d**, Percent of total *tfs1* reads derived from the *tfs1*^+^ (grey) and *tfs1*^*DN*^ alleles (black) in *pnmt1*+ repressible conditions (EMM 15uM thiamine) obtained from RNA-seq analysis of *epe1*∆ *tfs1*^*DN*^ and *epe1*∆ *tfs1*^+^ strains (no reads corresponding to the *tfs1*^*DN*^ allele were observed in the *epe1*∆ *tfs1*^+^ strain).

**Extended Data Figure 7.**
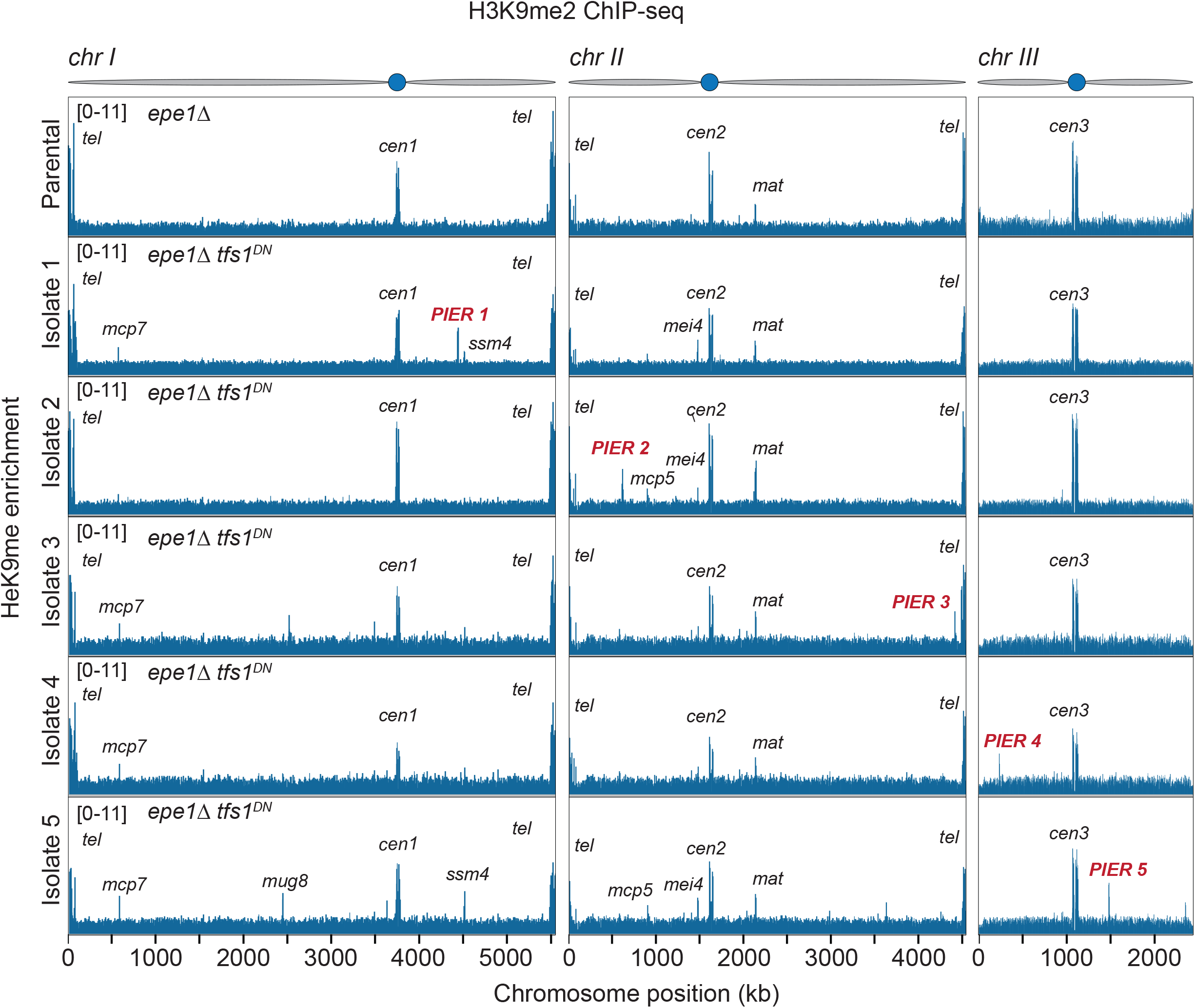
TFIIS^DN^ expression induces ectopic heterochromatin. Whole genome plots of H3K9me2 ChIP-seq enrichment for parental (*epe1*∆) and *epe1*∆ *tfs1*^*DN*^ isolates 1 through 5. Locations of PIERs noted in each plot.

**Extended Data Figure 8.**
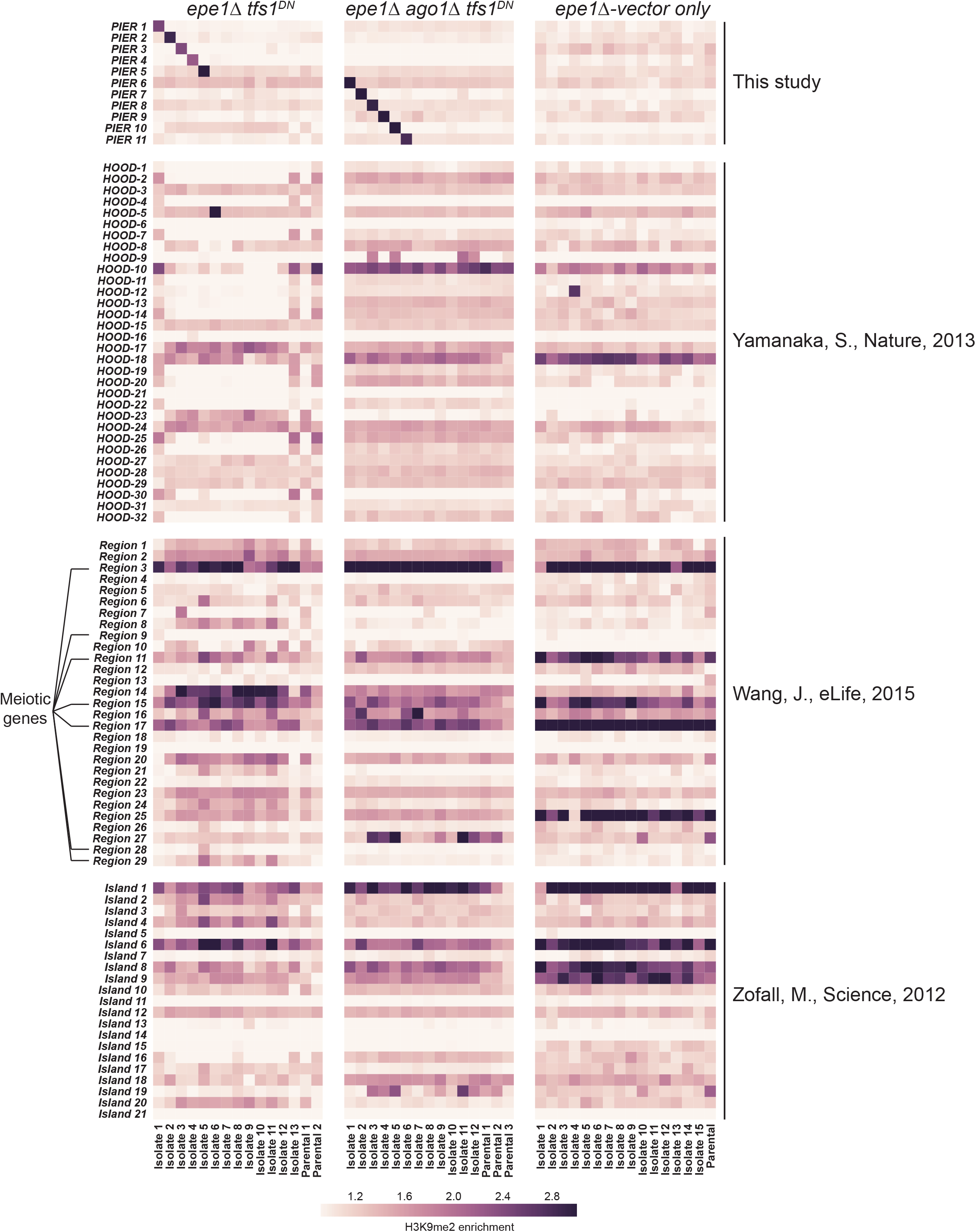
H3K9me enrichment at known heterochromatin nucleation sites and PIERs for all strains. PIERs and known sites of heterochromatin nucleation were analyzed for H3K9me enrichment (see Methods). All isolates and parental strains for each genotype are depicted. HOODs, H3K9me islands, and meiotic genes are indicated.

**Extended Data Figure 9.**
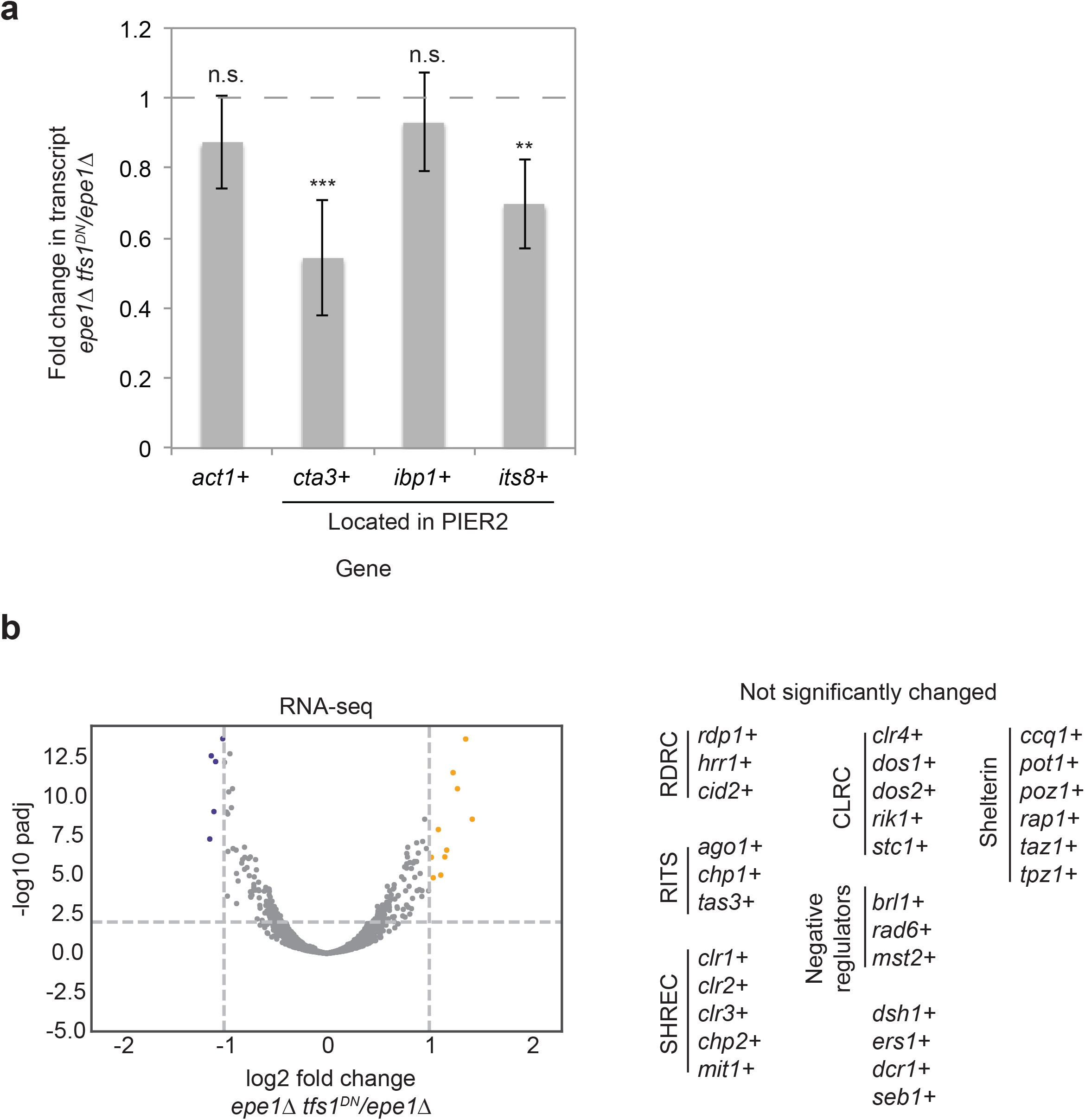
TFIIS^DN^ expression silences genes within a PIER but does not alter expression of heterochromatin factors. **a**, Significantly decreased transcript levels observed in two genes (*cta3*^+^ and *its8*^+^, but not *ibp1*^+^) found within PIER2 (*act1*^+^ is a control for no significant change). ***p<0.001, **p=0.0014. Dotted line = no change. **b**, RNA-seq volcano plot of log2 fold changes in transcript levels in *epe1*∆ *tfs1*^*DN*^ (Isolate 2 – containing PIER2) compared to *epe1*∆. 11 genes have significantly lower expression in *epe1*∆ *tfs1*^*DN*^ (blue dots); 15 genes have significantly higher expression in *epe1*∆ *tfs1*^*DN*^ (orange dots). padj = adjusted p-value obtained from DESeq2. (right) No significant changes in transcript levels were observed in heterochromatin factors.

**Extended Data Figure 10.**
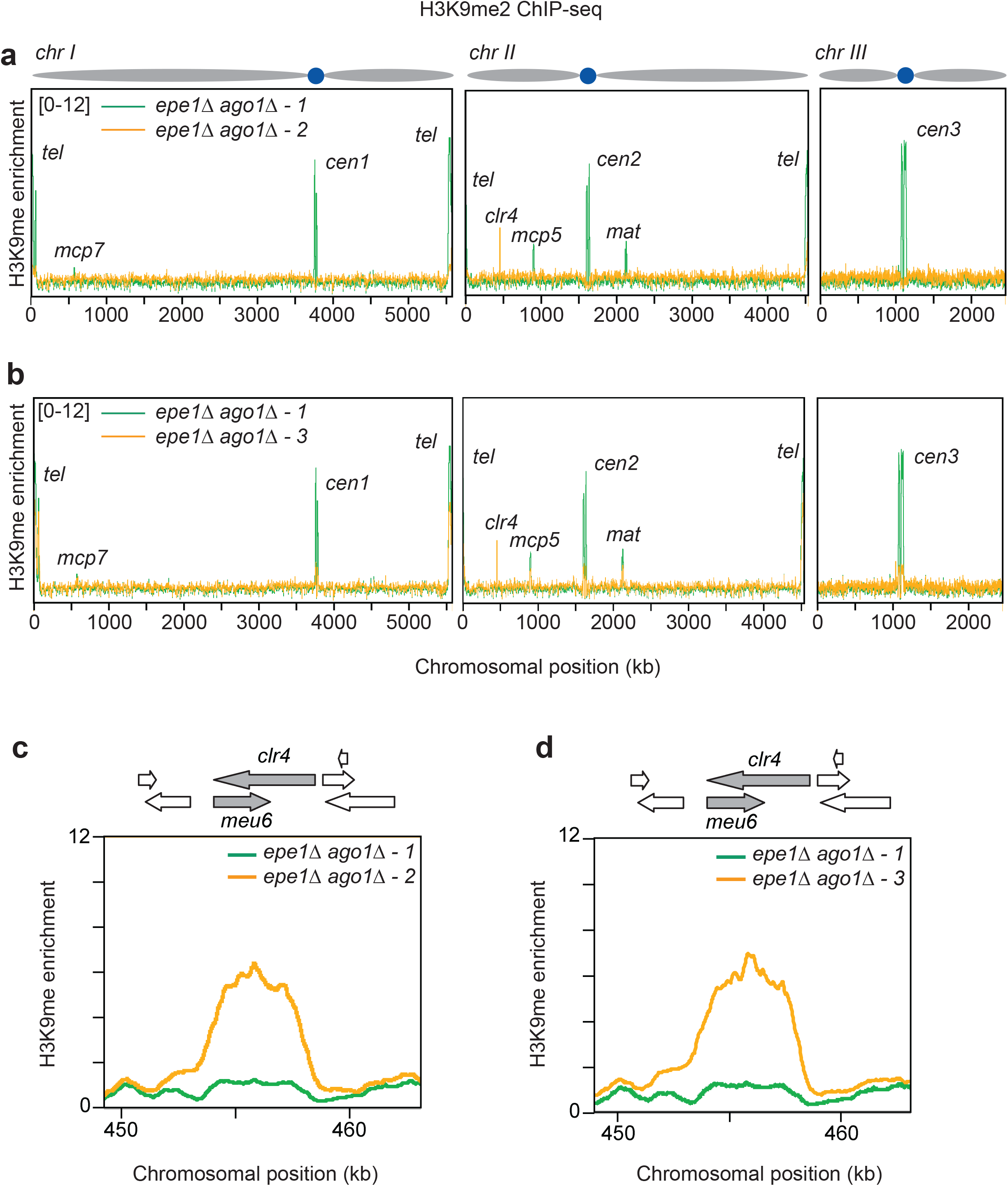
*epe1∆ ago1∆* strains can adaptively silence the *clr4*^+^ locus. **a**-**b**, Whole genome H3K9me2 enrichment plots for two independent *epe1*∆ *ago1*∆ strains that accumulated H3K9me2 over the *clr4*^+^ locus, *epe1*∆ *ago1*∆-*2* (**a**) and *epe1*∆ *ago1*∆-*3* (**b**) (in orange), compared to a isolate with normal distribution of H3K9me2 across the genome, *epe1*∆ *ago1*∆-*1* (in green). **c-d,** Genome browser image of the enrichment of H3K9me2 over *clr4*^+^ locus in *epe1*∆ *ago1*∆-*2* (**c**) and *epe1*∆ *ago1*∆-*3* (**d**).

**Extended Data Figure 11.**
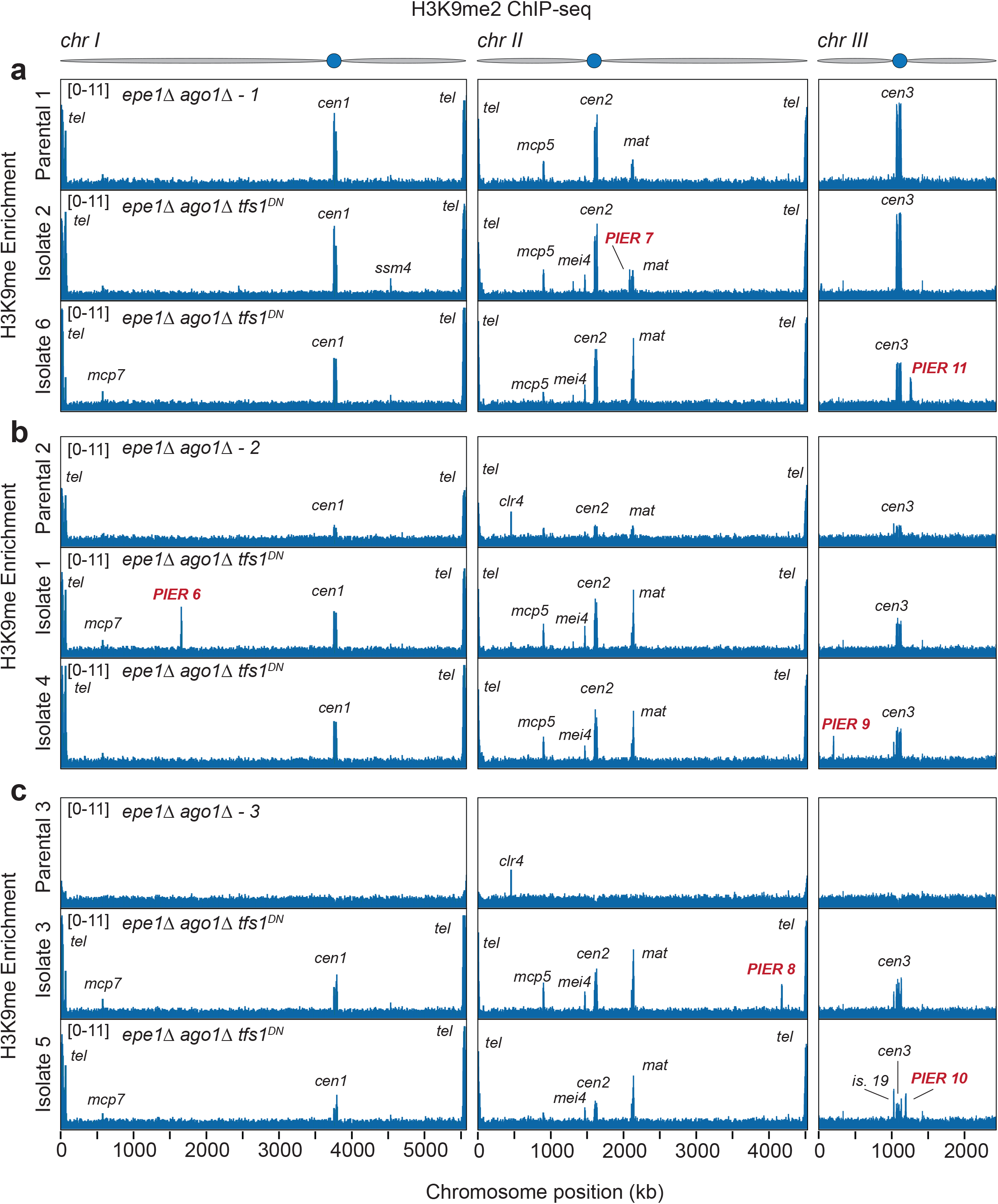
PIER formation is RNAi-independent and expression of TFIIS^DN^ can rescue the adaptive silencing of *clr4*^+^ observed in *epe1∆ ago1∆* strains. **a-c,** Genome wide plots of H3K9me2 enrichment. **a**, Parental (*epe1*∆ *ago1*∆-*1*) and two independent isolates of *epe1*∆ *ago1*∆ *tfs1*^*DN*^ derived from this parental are plotted depicting PIERs 7 and 11. **b**, Parental (*epe1∆ ago1∆-2*), initially harboring H3K9me2 at *clr4*^+^, and two independent isolates of *epe1*∆ *ago1*∆ *tfs1*^*DN*^ derived from this parental are plotted depicting PIERs 6 and 9. **c**, Parental (*epe1*∆ *ago1*∆-*3*), initially harboring H3K9me2 at *clr4*^+^, and two independent isolates of *epe1*∆ *ago1*∆ *tfs1*^*DN*^ derived from this parental are plotted depicting PIERs 8 and 10.

**Extended Data Figure 12.**
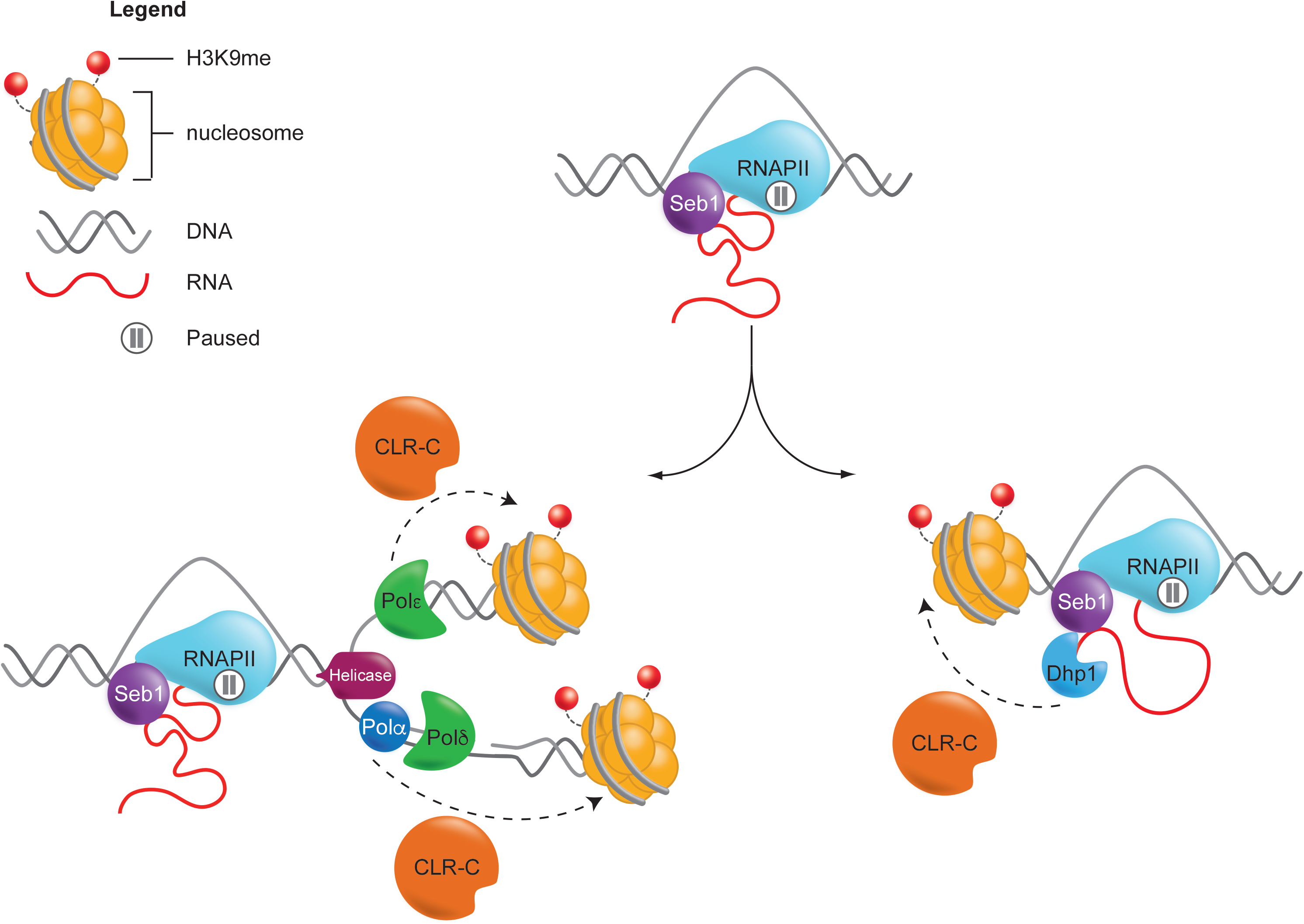
Speculative model for heterochromatin assembly mediated by Seb1-induced RNAPII Pausing. Engagement of Seb1 to RNA and RNAPII results in pausing of the RNAPII. Seb1-triggered RNAPII pausing may drive heterochromatin assembly by promoting the stalling of replisomes associated with CLR-C, via Polα or Polε, through RNAPII-replisome collisions. Seb1-induced pausing may also promote heterochromatin assembly via premature transcription termination and recruitment of Dhp1. RNAPII, RNA Polymerase II complex; CLR-C, Clr4 H3K9 methyltransferase complex; Polδ, DNA polymerase delta; Polα, DNA polymerase alpha; Polε, DNA polymerase epsilon.

